# The Effect on Speech-in-Noise Perception of Real Faces and Synthetic Faces Generated with either Deep Neural Networks or the Facial Action Coding System

**DOI:** 10.1101/2024.02.05.578468

**Authors:** Yingjia Yu, Anastasia Lado, Yue Zhang, John F. Magnotti, Michael S. Beauchamp

**Affiliations:** Department of Neurosurgery, Perelman School of Medicine, University of Pennsylvania; Department of Neurosurgery, Baylor College of Medicine

## Abstract

The prevalence of synthetic talking faces in both commercial and academic environments is increasing as the technology to generate them grows more powerful and available. While it has long been known that seeing the face of the talker improves human perception of speech-in-noise, recent studies have shown that synthetic talking faces generated by deep neural networks (DNNs) are also able to improve human perception of speech-in-noise. However, in previous studies the benefit provided by DNN synthetic faces was only about half that of real human talkers. We sought to determine whether synthetic talking faces generated by an alternative method would provide a greater perceptual benefit. The facial action coding system (FACS) is a comprehensive system for measuring visually discernible facial movements. Because the action units that comprise FACS are linked to specific muscle groups, synthetic talking faces generated by FACS might have greater verisimilitude than DNN synthetic faces which do not reference an explicit model of the facial musculature. We tested the ability of human observers to identity speech-in-noise accompanied by a blank screen; the real face of the talker; and synthetic talking face generated either by DNN or FACS. We replicated previous findings of a large benefit for seeing the face of a real talker for speech-in-noise perception and a smaller benefit for DNN synthetic faces. FACS faces also improved perception, but only to the same degree as DNN faces. Analysis at the phoneme level showed that the performance of DNN and FACS faces was particularly poor for phonemes that involve interactions between the teeth and lips, such as /f/, /v/, and /th/. Inspection of single video frames revealed that the characteristic visual features for these phonemes were weak or absent in synthetic faces. Modeling the real *vs.* synthetic difference showed that increasing the realism of a few phonemes could substantially increase the overall perceptual benefit of synthetic faces, providing a roadmap for improving communication in this rapidly developing domain.

## Introduction

Recent advances in computer graphics have made it much easier to create realistic, synthetic talking faces, spurring adoption in commercial and academic communities. For companies, agents created by pairing a synthetic talking face with the output of large language models provide an always-available simulacrum of a real human representative (Perry et al., 2023). In academia, the use of synthetic talking faces in studies of speech perception provides more precise control over the visual features of experimental stimuli than is possible with videos of real human talkers (Thézé et al., 2020).

Of particular interest is the long-standing observation that humans understand speech-in-noise much better when it is paired with a video of the talker’s face (Sumby and Pollack, 1954). The ability to rapidly generate a synthetic face saying arbitrary words suggests the possibility of an “audiovisual hearing aid” that displays a synthetic talking face to improve comprehension. This possibility received support from two recent studies that used deep neural networks (DNNs) to generate realistic, synthetic talking faces (Shan et al., 2022; Varano et al., 2022). Both studies found that viewing synthetic faces significantly improved speech-in-noise perception, but the benefit was only about half as much as viewing a real human talker.

The substantial disadvantage of synthetic faces raises the question of whether alternative techniques for generating synthetic faces might provide a greater perceptual benefit. DNNs associate given speech sounds with visual features in their training dataset, but do not contain any explicit models of the facial musculature. In contrast, the facial action coding system (FACS) uses 46 basic action units to represent all possible movements of the facial musculature that are visually discernable (Ekman and Friesen, 1978, 1976; Parke and Waters, 2008). Unlike DNNs, the FACS scheme is built on an understanding of the physical relationship between speech and facial anatomy, potentially resulting in more accurate representations of speech movements. To test this idea, we undertook a behavioral study to compare the perception of speech-in-noise on its own; speech-in-noise with real faces (to serve as a benchmark); and speech-in-noise presented with two types of synthetic faces. The first synthetic face type was generated by a deep neural network, as in the studies of (Shan et al., 2022; Varano et al., 2022). The second synthetic face type was generated using FACS, as implemented in the commercial software package JALI (Edwards et al., 2016; Zhou et al., 2018). For comparison with previous studies, we performed a word-level analysis in which each response was scored as correct or incorrect. To facilitate more fine-grained comparisons between the different face types, we also analyzed data using the phonemic content of each stimulus word.

## Methods

### Participant recruitment and testing

All experiments were approved by the Institutional Review Board of the University of Pennsylvania, Philadelphia, PA. Participants were recruited and tested using Amazon Mechanical Turk (https://www.mturk.com/), an online platform that provides access to an on-demand workforce. Only “master workers” were recruited, classified as such by Amazon based on their high performance and location in the U.S.A. Sixty-two master workers completed the main experiment (median time to complete: 12 minutes) and received $5 reimbursement. The workers answered the questions “Do you have a hearing impairment that would make it difficult to understand words embedded in background noise?” and “Do you have an uncorrected vision impairment that would make it difficult to watch a video of a person talking?” One worker was excluded because of a reported hearing impairment, leaving sixty-one participants whose data is reported here. There were 25 females and 36 males, mean age 46, range 30 to 72.

### Overview

Sixty-one participants identified seventy-three words presented in five different formats (Figure 1). Sixty-four of the words contained added auditory noise to make identifying them more difficult and increase the importance of visual speech. There were four formats of noisy words: auditory-only (An); with a talking face (audiovisual; AnV) that was either the real face of the talker (AnV:*Real*); a synthetic face created using the facial action coding system (AnV:*FACS*); or a synthetic face created using a deep neural network (AnV:*DNN*). To prevent perceptual learning, each word was only presented once to each participant, 16 words in each of the four formats. Within participants, the order of words and face formats was randomized, and across participants, the format of each word was cycled to ensure that every word was presented in every format. To assess participant compliance, the remaining nine words presented were clear audiovisual words (AV:*catch_trials*). The catch trials sampled all face types (3 Real, 3 FACS, 3 DNN) and the talkers and words differed from those presented in the noisy trials to prevent learning. Accuracy for catch trials was very high (mean of 98%) demonstrating attention and task engagement. All data was analyzed in R, primarily using mixed effects models. See *Supplementary Materials* for all data and an R markdown document that contains all analysis code and results.

**Figure 1.**
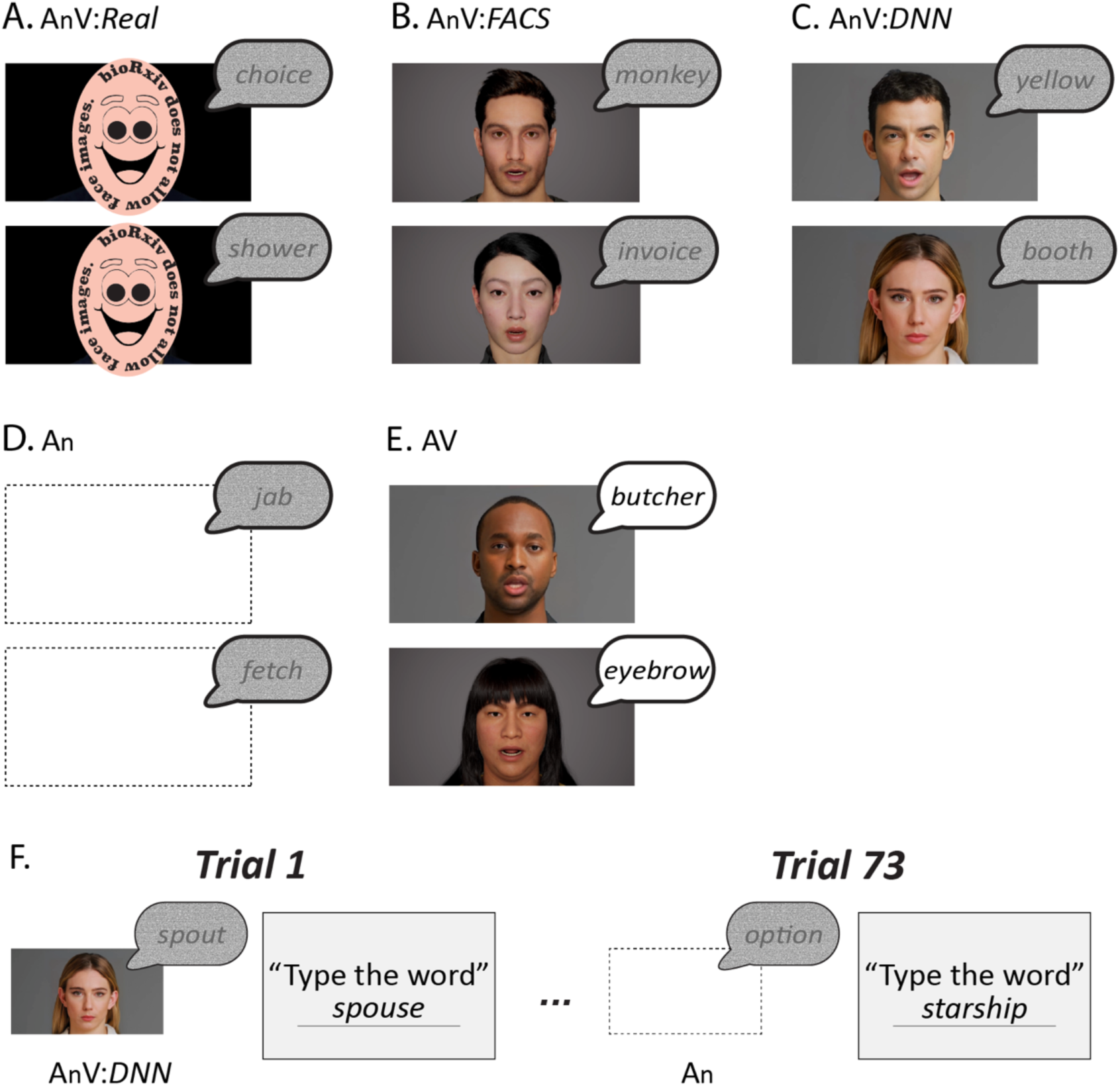
**A.** The main stimulus set consisted of 64 single words with auditory noise added, 32 recorded by a male talker and 32 recorded by a female talker. The noisy words were presented in four formats. The first format consisted of the noisy auditory recordings paired with a video of the actual talker (AnV:*Real*). **B.** The second format consisted of the recordings paired with gender-matched synthetic face movies generated by a facial action coding system model (AnV:*FACS*). **C.** The third format consisted of the recordings paired with synthetic face movies generated by a deep neural network (AnV:*DNN*). The gender of the synthetic face matched the gender of the voice. **D.** The fourth format consisted of the recordings presented with a blank screen (An). Note that for (A)-(D), the auditory component of the stimulus was identical, only the visual component differed. **E.** In catch trials, an additional stimulus set was presented consisting of recordings of 9 audiovisual words without added noise (AV) paired with gender-matched real, FACS, or DNN faces (three words each). The words, faces and voices were different than in the main stimulus set to prevent any interference. **F.** Each participant was presented with 73 words (64 noisy and 9 clear) in random order. Within participants, each noisy word from the main stimulus set was presented only once, in one of the four formats. Counterbalancing was used to present every noisy word in each of the four formats shown in (A)-(D). For instance, for participant 1, the word *spout* was presented in AnV:*DNN* format, while for participant 2, *spout* was presented in AnV:*Real* format, *etc.* Following the presentation of each word, participants typed the word in a text box.

### Subject responses and scoring: Word-level

Following presentation of a word, participants were instructed to “type the word” into a text box; the next trial did not begin until a response was entered. If a participant’s response matched the stimulus word, the trial was scored as “correct”, otherwise the trial was scored as “incorrect”. For example, the stimulus word *wormhole* and the response *wormhole* was correct, while the stimulus word *booth* and the response *boot* was incorrect. Misspellings were not considered incorrect (*e.g.* stimulus *echos* and response *echoes* was correct) nor were homophones (*e.g.* stimulus *wore* and response *war* was correct.) The analysis was performed separately for each condition (An, AnV:*Real*, AnV:*DNN*, AnV:*FACS*, AV:*catch_trials*). Mean accuracy for each condition was calculated per participant, and then averaged across participants. See *Supplementary Materials* for a complete list of the words, response and scores.

### Phonemic analysis

In addition to the binary word-level accuracy measure, a continuous accuracy measure was calculated for each trial based on the overlap in the phonemes in the stimulus word and the response. Phoneme composition was determined using the Carnegie Mellon University (CMU) pronouncing dictionary (http://www.speech.cs.cmu.edu/cgi-bin/cmudict). Lexical stress markers were removed from vowels. For responses that contained multiple words, the phonemes for all response words were extracted from the CMU dictionary and entered into the calculation. Repetitions of the same phoneme were also entered into the calculation. The accuracy measure was the Jaccard index: the number of phonemes in common between the stimulus and response divided by the total number of phonemes in the stimulus and response. The measure ranged from 0 (no phonemes in common between stimulus and response) to 1 (identical phonemes in stimulus and response) and was calculated as

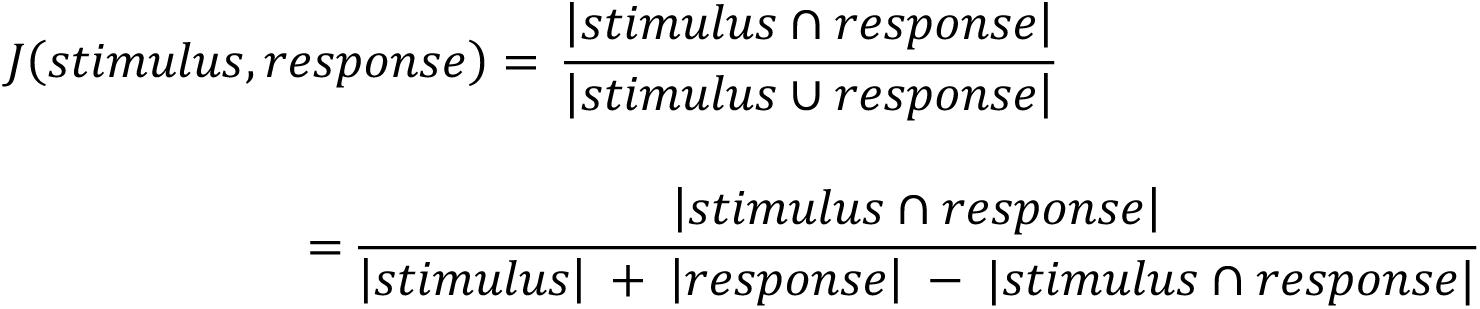

For example, the stimulus word *polish* contains the phonemes P, AA, L, IH, SH. One participant’s response was *policy*, containing the phonemes P, AA, L, AH, S, IY. The intersection contains 3 phonemes (P, AA, L) while the union contains 8 phonemes (P, AA, L, IH, SH, AH, S, IY) for a Jaccard index of 3/8 = 38%. In another example, the stimulus word *ethic* contains the phonemes EH, TH, IH, K while the response *essay* contains the phonemes EH, S, EY. The intersection contained 1 phoneme and the union contained 6 phonemes, for a Jaccard index 1/6 = 17%. See *Supplementary Materials* for a complete list of Jaccard indices.

The analysis was performed separately for each stimulus condition. For participant-level analysis, the mean accuracy across all trials was calculated for each participant, and then averaged across participants. For phoneme-specific analysis, the number of times the phoneme was successfully identified was divided by the total number of times the phoneme was presented across all words, across all participants.

### Additional stimulus details

The original stimulus material consisted of 32 audiovisual words recorded by a female talker and 32 words recorded by a male talker. Pink noise was added to the auditory track of each recording at a signal-to-noise ratio (SNR) of -12dB. For the An format, only the noisy auditory recording was played with no visual stimulus. For the AnV:*Real* format, the audio recording was accompanied by the original video recording. For the synthetic faces, gender-matched synthetic faces roughly approximating the appearance of the real talkers were created. During the online testing procedure, videos were presented using a custom JavaScript routine that ensured that all stimuli were presented with the same dimensions (height: 490 pixels; width: 872 pixels) regardless of the participant’s device.

AnV:*FACS* words were created using JALI software (Edwards et al., 2016)(https://jaliresearch.com/). The text transcript was manually tuned to create more pronounced mouth movements (*e.g.*, AWLTHOH for although; see *Supplementary Materials* for a complete list of the phonetic spelling). Following JALI animation, the mouth movements were manually adjusted with MAYA’s graph editor to better match the mouth movements in the real videos. The edited animation sequence was imported into Unreal Engine 5.10 and rendered as 16:9 images at 50 mm focal length and anti-aliasing with 16 spatial sample count. Image sequences were assembled into mp4 format and aligned with the original audio track in Adobe Premiere. The video frame rates was 24 fps.

AnV:*DNN* words were created with D-ID studio (https://www.d-id.com/). Two artificial faces included with D-ID studio were used as the base face. Clear audio files from the real speakers were uploaded, and the static driver-3 was used to minimize head and neck movements. Eye blinking and watermarks in each movie were removed using Adobe Premiere and exported in mp4 movie at 24 fps.

## Results

### Participant-level analysis: Word accuracy

In the first analysis, responses were scored as correct if they exactly matched the stimulus word and incorrect otherwise. Seeing the face of the talker improved the intelligibility of noisy auditory words. For real faces, accuracy increased from 10% in the auditory-only condition (An) to 59% in the audiovisual condition (AnV:*Real*), averaged across words and participants. There was also an improvement, albeit smaller, for synthetic faces (*Figure 2A)*. From the auditory-only baseline of 10%, accuracy improved to 29% with faces generated by the facial action coding system (AnV: *FACS*). Accuracy was 30% for faces generated by a deep neural network (AnV:*DNN*). While there was a range of accuracies across participants, accuracy was higher for the real face format than the auditory-only format in 60 of 61 participants and higher for real faces than synthetic faces (*Synthetic;* average across *DNN* and *FACS*) in 59 of 61 participants (*Figure 2B*).

**Figure 2.**
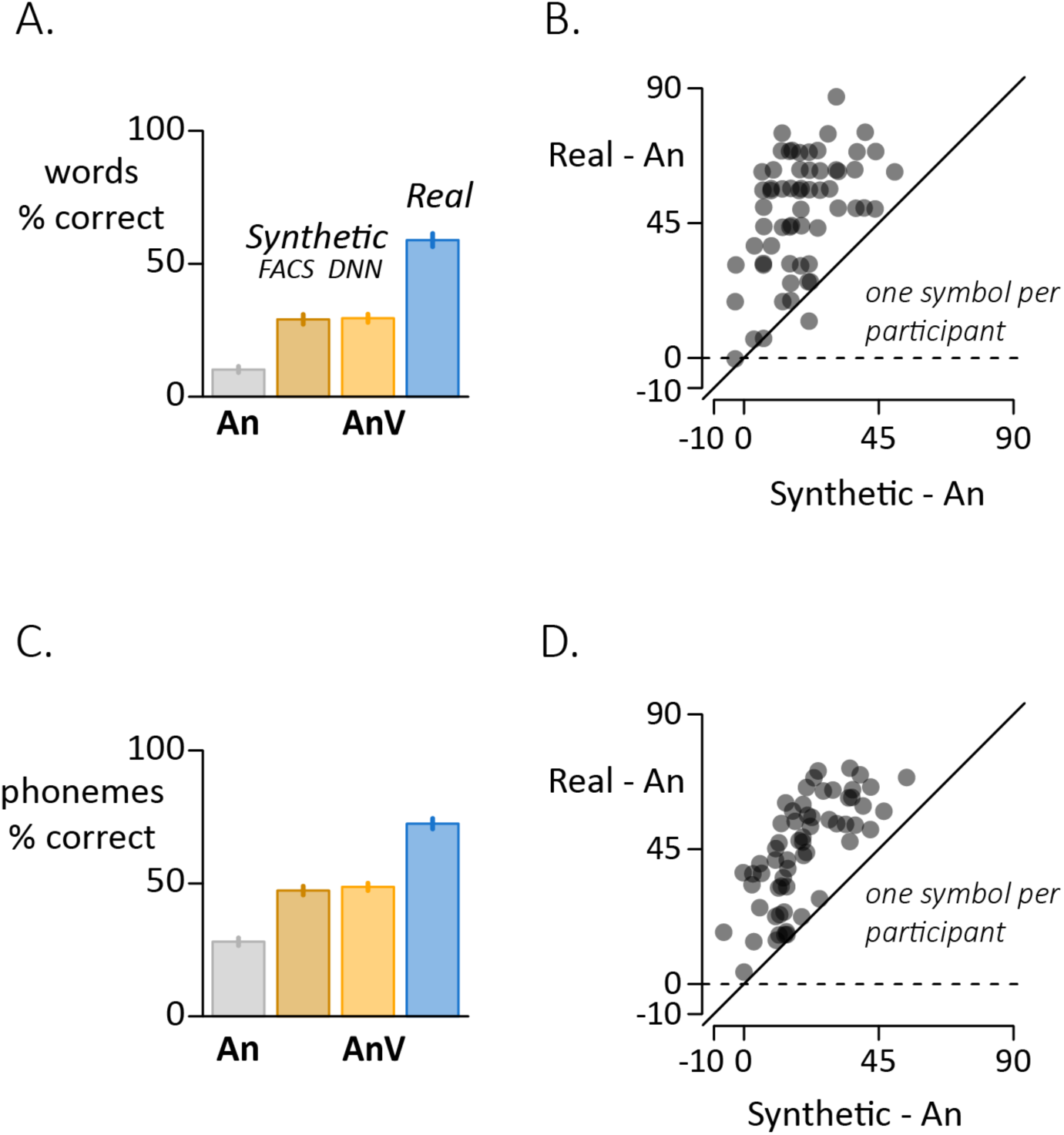
**A.** For word-level scoring, the response was assessed as correct if it exactly matched the stimulus word and incorrect if it did not. Each bar shows the mean accuracy for each stimulus format (error bars show the standard error of the mean across participants). B. Variability across participants assessed with word-level scoring (one symbol per participant). The y-axis shows the perceptual benefit of real faces (real minus auditory-only accuracy). The x-axis shows the perceptual benefit of synthetic faces (average of *FACS* and *DNN* accuracies minus auditory-only). Participants above the dashed line show a benefit for real faces compared with auditory-only. Participants above the solid identity line show a greater benefit for real faces than synthetic faces. **C.** For phoneme-level scoring, the phonemic content of the stimulus and response were compared and the percentage of correct phonemes calculated for each stimulus format. **D.** Variability across participants assessed with phoneme-level scoring (one symbol per participant).

To estimate the statistical significance of these observations, a mixed-effects model was constructed with a dependent variable of accuracy; fixed effect of word format; and random effects of word, participant and participant batch (complete model specification and output in *Supplementary Materials*). There was a main effect of stimulus format (ξ^2^*3* = 768, *p* < 10^-16^) and *post hoc* pair-wise comparisons showed that words accompanied by a visual face (real or synthetic) were perceived more accurately than auditory-only words (all *p* < 10^-16^). The accuracy for real faces was higher than for synthetic faces (*Real vs. DNN*; *t* = −17, *p* < 10^-16^; *Real vs. FACS*; t = −17, *p* < 10^-16^), but was equivalent for the two synthetic face formats (*t* = 0.2, *p* = 0.99).

### Participant-level analysis: Phoneme accuracy

In a second analysis, instead of classifying each response as either correct or incorrect, partial credit was given if the response contained phonemes that matched those in the stimulus word. This scoring method generated a phoneme accuracy score for each condition in each participant. The pattern of results were very similar to the word accuracy analysis (*Figure 2C*; *Supplementary Materials*). There was a main effect of stimulus format (ξ^2^*3* = 986, *p* < 10^-16^) and *post hoc* pair-wise comparisons showed that words accompanied by a visual face (real or synthetic) were perceived more accurately than auditory-only words (all *p* < 10^-16^). The accuracy for real faces was higher than for synthetic faces (*Real vs. DNN*; *t* = −17, *p* < 10^-16^; *Real vs. FACS*; t = −18, *p* < 10^-16^) with equivalent accuracy for the two synthetic face formats (*t* = 0.9, *p* = 0.82). Accuracy was higher for real faces than synthetic faces in every participant (*Figure 2D*).

### Phoneme-level analysis

In a third analysis, the accuracy difference between real and synthetic faces was examined separately for each phoneme (*Figure 3A*). For four phonemes (*/th/*, */dh/*, */f/*, */v/*) the accuracy advantage of real faces was especially pronounced (*Real* >> *Synthetic*). This observation could arise because these phonemes had particularly high accuracy for the *Real* format, particularly low *Synthetic* accuracy, or both. To distinguish these possibilities, we calculated the mean *Real* accuracy for (*/th/*, */dh/*, */f/*, */v/*) compared with other phonemes, and found little difference (78% *vs.* 78%, *t* = 0.1, *p* = 0.96, unequal variance, unpaired *t*-test). In contrast, the mean *Synthetic* accuracy was significantly lower for (*/th/*, */dh/*, */f/*, */v/*) than for other phonemes (29% *vs.* 62%, *t* = −5, *p* = 0.004). Thus, the greater real−synthetic difference for */th/*, */dh/*, */f/*, */v/* than other phonemes (48% *vs.* 17%, *t* = 11, *p* = 10^-4^) was attributable to particularly low *Synthetic* accuracy (*Figure 3B*).

**Figure 3.**
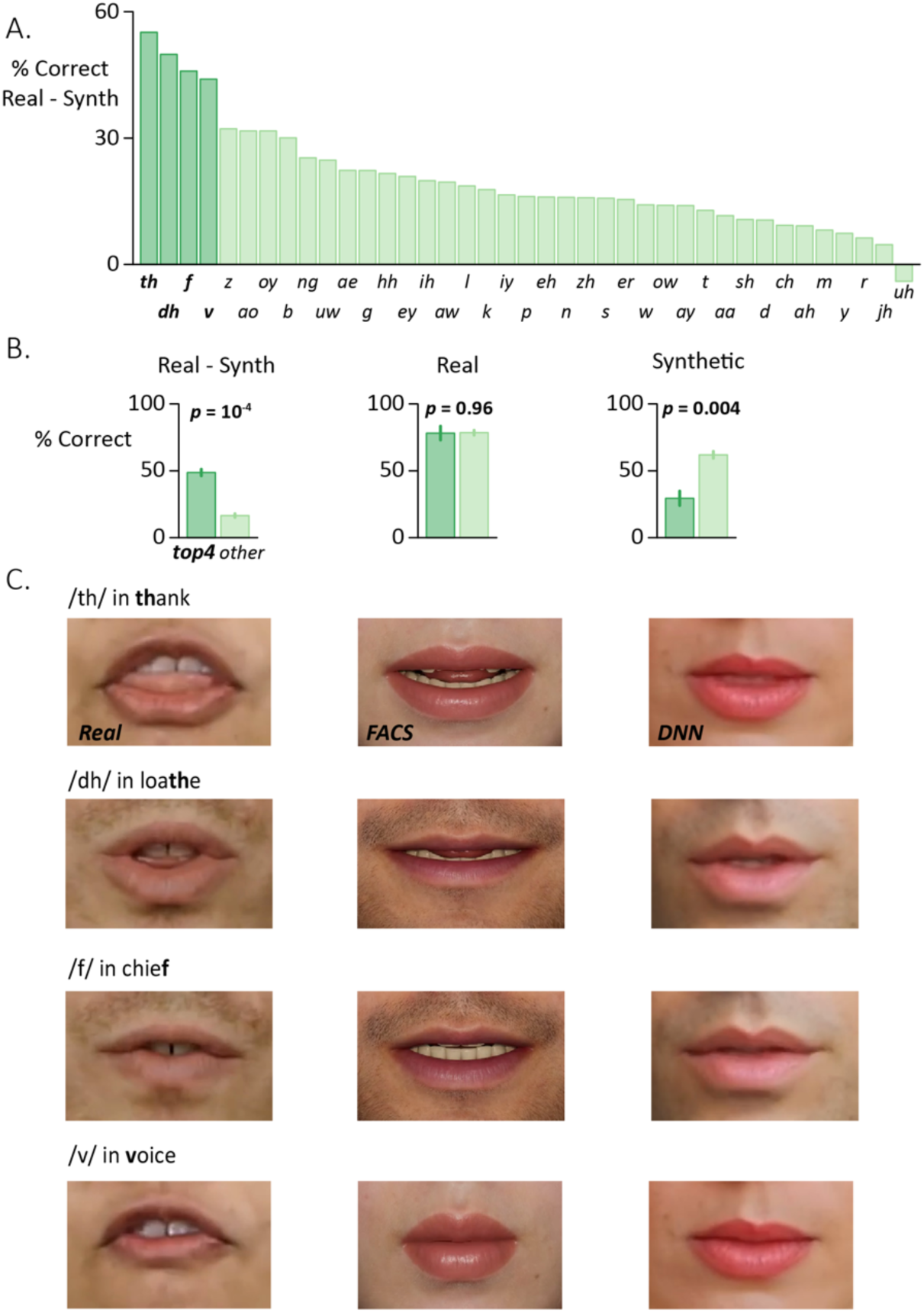
**A.** For each phoneme, the perceptual accuracy was calculated separately for each stimulus format across all participants. The accuracy for audiovisual synthetic faces (average of *DNN* and *FACS*) was subtracted from the accuracy for audiovisual real faces to generate a single value for each phoneme. Phonemes were sorted by real−synthetic difference to highlight four phonemes (dark green bars) that showed a larger real−synthetic difference than the other phonemes (light green bars). **B.** Four phonemes (*/th/*, */dh/*, */v/*, */f/*) showed a significantly larger real−synthetic difference than the other phonemes (left plot). Average of these four phonemes shown by dark green bar, average of other phonemes shown by light green bar, error bar shows SEM. This was not due to differences in the real face condition (middle plot) but rather to low accuracy for the top four phonemes in the synthetic face condition (right plot). **C.** Enlargement of the mouth region for the three different face formats for words containing (*/th/*, */dh/*, */v/*, */f/*). Enlargement for illustration only, participants viewed the entire face, as shown in Figure 1. Top row: video frame 18 from the word *thank*. Second row: video frame 30 from the word *loathe*. Third row: video frame 27 from the word *chief*. Fourth row: Video frame 16 from the word *voice*.

The poor accuracy for synthetic */th/*, */dh/*, */f/*, */v/* suggested that some key visual features might be missing (*Figure 3C*). For */th/* and */dh/*, the salient visual feature is the tongue sandwiched between the teeth. This feature was clearly visible in the real face videos but was absent from the *DNN* and *FACS* face videos. For */f/* and */v/*, the salient visual feature is the upper teeth pressed onto the lower lip. This feature was obvious in the real face videos but not in the synthetic face videos.

### Modeling the effects of improving the four phonemes

With improvements in computer graphics, it should be possible to create synthetic faces that more accurately depict */th/*, */dh/*, */f/*, */v/* and thereby increase the synthetic face benefit for words containing these phonemes. To estimate this increase, we constructed a logistic model that predicted the word accuracy based on the presence or absence of every different phoneme in the word. The fitted model contained one coefficient for each synthetic face phone and one coefficient for each real face phoneme, and was a good fit to the data (*r*^2^ = 0.85, *p* < 10^-16^). In the hypothetical best case, the improved synthetic versions of */th/*, */dh/*, */f/*, */v/* would be as good as the real face versions. This was simulated in the model by replacing the synthetic face coefficients for these phonemes with the real face coefficients. With this adjustment, the model predicted a word accuracy of 43%, compared with 29% for the original versions of */th/*, */dh/*, */f/*, */v/*, suggesting that improving the quality of the synthetic faces for the four phonemes with very low synthetic benefit could significantly boost overall accuracy.

### Effects of alternative stimulus material

The overall phoneme accuracy rates shown in *Figure 2C* were determined by the phonemic content of the 64 tested words. This raises the question of how the results might differ for a much larger corpus of words. With a large enough corpus, the frequency of phonemes should match their overall prevalence in the English language. To simulate an experiment with a large corpus, we weighted the real−synthetic difference for each phoneme by its prevalence in the English language (Hayden, 1950). This procedure predicted an overall auditory-only phoneme accuracy of 40%, compared with 37% for the actual stimulus set. The predicted synthetic face phoneme accuracy was 60%, compared with 58% for the actual stimulus set, and the predicted real face accuracy was 78%, compared with 79% for the tested words. The predicted real−synthetic difference was 18% compared with 21% for the actual stimulus set. In short, testing with a large corpus of words (or a smaller set of words that matched overall English-language phoneme frequency) should lead to only small changes in accuracy.

## Discussion

Our study replicates decades of research by showing that seeing the face of a real talker improves speech-in-noise perception (Peelle and Sommers, 2015; Sumby and Pollack, 1954). Our study also confirms two recent reports that viewing a synthetic face generated by a deep neural network (DNN) significantly improves speech-in-noise perception (Shan et al., 2022; Varano et al., 2022). Both the present study and these previous reports found that the improvement from viewing DNN faces was only about half that provided by viewing real faces.

To determine if some idiosyncrasy of DNN faces was responsible for their poor performance relative to real faces, we also tested synthetic faces generated with a completely different technique, the facial action coding system (FACS). Since FACS explicitly models the relationship between the facial musculature and visual speech movements, we anticipated that it might provide more benefit to speech perception than DNN faces. Instead, the perceptual benefit of FACS faces was very similar to that of DNN faces.

To better understand the real−synthetic difference, we decomposed stimulus words and participant responses into their component phonemes. This analysis revealed variability across phonemes. Four phonemes (*/th/*, */dh/*, */f/*, */v/*) had an especially large real−synthetic difference, driven by low performance for both DNN and FACS faces.

Examining single frames of the real and synthetic videos for words containing these phonemes revealed an obvious cause for the reduced benefit of synthetic faces. The synthetic videos were missing the interactions between teeth, lip and tongue that are the diagnostic visual feature for */th/*, */dh/*, */f/*, */v/*. Without these visual cues, participants did not receive the visual information beneficial for detecting these phonemes in speech-in-noise.

### The concept of visemes

Visemes can be defined as the set of mouth configurations used to pronounce the phonemes in a language and may be shared across different phonemes. For instance, (*/f/*, */v/*) are visually similar labiodental fricatives that require talkers to place the top teeth on the lower lip. The acoustic difference is generated by voicing (*/f/* is unvoiced while */v/* is voiced); this voicing difference does not provide a visible cue to an observer. Similarly, (*/th/*, */dh/*) are dental fricatives, articulated with the tongue against the upper teeth, with */th/* unvoiced and */dh/* voiced. While there is no generally agreed upon set of English visemes, five common viseme classifications all place (*/f/*, */v/*) in one viseme category and (*/th/*, */dh/*) in a different viseme category (Cappelletta and Harte, 2012). Our results confirm the veracity of this grouping.

### How to improve synthetic faces

For the FACS faces, it would be possible to manually control teeth and tongue positioning using the underlying 3D face models, although this would be a time-consuming process. Alternately, the automated software used to animate the 3D face models (JALI)(Edwards et al., 2016) could be modified to automatically code teeth and tongue positioning. For DNN faces, it is less straightforward to incorporate dental and labial interactions. The DNN models are trained on thousands or millions of examples of auditory and visual speech, and the network learns the correspondence between particular sounds and visual features. It may be that dental and labial interactions are highly variable across talkers, or not easily visible in the videos used for training, resulting in their absence in the final output. A common step in neural network model creation is fine-tuning. Incorporating training data that explicitly includes dental and labial features, such as from electromagnetic articulography (Schönle et al., 1987) or MRI (Baer et al., 1987), would improve the DNN’s ability to depict these features.

### Other factors

While four phonemes showed an especially large real−synthetic difference (mean of 48% for */th/*, */dh/*, */f/*, */v/*) there was also a substantial difference for the remaining phonemes (mean of 16%). The origin of this difference is likely to be multi-faceted. One likely contributing factor is that just as for */th/*, */dh/*, */f/*, */v/*, the diagnostic mouth features for other phonemes are not as accurately depicted or as obvious in the synthetic faces as they are in the real face videos. This possibility could be tested by showing the most diagnostic single frame from each type of video and asking participants to guess the phoneme being spoken. The prediction is that, even for single video frames, synthetic performance would be worse than real face performance. Differences between DNN faces and real faces when pronouncing particular phonemes has been proposed as a method to detect deep-fake videos (Agarwal et al., 2020) although the phonemes examined in this study (*/m/*, */b/*, */p/*) were not those that showed the largest real−synthetic difference inthe present study. For real talkers, information about speech content is available throughout the face. Since synthetic face generation concentrates on the mouth and lip region, this decreases the information about speech content available to observers (Munhall and Vatikiotis-Bateson, 2004).

Another contributing factor for the real−synthetic difference could be temporal synchrony. The alignment between auditory and visual speech contributed to causal inference and other perceptual processing underlying speech perception. While the DNN and FACS synthetic faces were aligned to the auditory speech, this alignment will not be as precise as the alignment with the real talker from which the auditory and visual speech were recorded. To assess the contribution of this factor, several approaches are possible. One would be to use voice and face from two different talkers (or a synthesized voice) so that any alignment imprecision is similar between the real and synthetic faces (Bhat et al., 2015; Magnotti et al., 2013).

### Relevance to experimental studies of audiovisual speech perception

An important reason for creating synthetic talking faces is to investigate audiovisual speech perception (Thézé et al., 2020). In the well-known illusion known as the McGurk effect, incongruent auditory and visual speech leads to unexpected percepts (McGurk and MacDonald, 1976). However, different McGurk stimuli vary widely in their efficacy. For instance, Basu Mallick and colleagues tested 12 different McGurk stimuli used in published studies, and found that the strongest evoked the illusion on 58% of trials while the weakest stimulus evoked the illusion on only 17% of trials (Basu Mallick et al., 2015). The source of such high levels of inter-stimulus variability is unclear and difficult to study experimentally, as talkers have only limited ability to consciously control different aspects of their speech production. In contrast, synthetic faces, especially those created with FACS and related techniques, provide the ability to precisely control every aspect of the visual speech (Thézé et al., 2020).

### Evidence from incongruent audiovisual speech

Theze and colleagues tested the incongruent pairing of a synthetic face pronouncing visual */v/* with an auditory */b/* auditory phoneme and found that it induced the percept of */v/* on about 40% of trials (Thézé et al., 2020). For comparison, the same pairing with a real visual face evoked the percept of */v/* on 94% of trials (Dias et al., 2016; Shahin, 2019). Taken together, this indicates that for incongruent auditory-visual speech, synthetic faces influenced perception much less than real faces, consistent with the real−synthetic difference for speech-in-noise observed in the present study and (Shan et al., 2022; Varano et al., 2022).

### Experimental predictions for incongruent visual speech

The data from (Thézé et al., 2020) and (Dias et al., 2016) on the incongruent visual */v/* and auditory */b/* pairing showed a large real−synthetic difference (54%; 94% *vs.* 40%). The original study of McGurk *et al*. tested the visual phonemes */g/* and */k/* (McGurk and MacDonald, 1976). In the present study of noisy speech, the phonemes */f/* or */v/* displayed an approximately threefold larger real−synthetic difference than */g/* and */k/*. Therefore, our results predict a smaller real−synthetic difference for McGurk syllables containing */g/* or */k/* than for incongruent audiovisual speech containing visual */v/* or */f/*. Since the real−synthetic difference for visual */v/* and auditory */b/* was 54%, then the real−synthetic difference for visual */g/* and auditory */b/* would be expected to be about a third as large (18%). For instance, if the McGurk pairing of auditory */b/* and a real face enunciating */g/* evoked the fusion percept of */d/* on 50% of trials, then the same pairing with a synthetic face should evoke the */d/* percept on about 32% of trials.

### Limitations of the present study

The present study has a number of limitations. In order to maximize the number of tested words and minimize experimental time, only a single noise level was tested, as in a previous study of DNN faces (Varano et al., 2022), with a high level of noise selected to maximize the benefit of visual speech (Rennig et al., 2020). Another previous study of DNN faces tested multiple noise levels and found a lawful relationship between different noise levels and perception (Shan et al., 2022). As the amount of added auditory noise decreased, accuracy increased for the no-face, real face and DNN face conditions in parallel, converging at ceiling accuracy for all three conditions when no auditory noise was added. We would expect a similar pattern if our experiments were repeated with different levels of auditory noise. While our study only examined speech perception, a similar approach could be taken to compare real and synthetic faces in other domains, such as emotions and looking behavior (Miller et al., 2023).

### Advantages of phonemic analysis

One relative novelty in our analysis was examining the contribution of different phonemes. While these results need to be validated in independent samples, it suggests that it should be possible to estimate the AV improvement or real or synthetic faces for an arbitrary set of words.

Phonemic analysis may also be helpful in untangling the sources of the substantial individual differences observed in the present study and other work on audiovisual speech perception (Grant et al., 1998; Rennig et al., 2020). Individual variability could be uniform across different phonemes, or it could be that some participants are especially poor at perceiving the visual speech linked to specific phonemes, in which case a training regimen could be targeted at just those phonemes.

## Conclusions

Massaro and Cohen pioneered the use of synthetic faces to examine audiovisual speech perception (Massaro and Cohen, 1995) and recent advances in computer graphics and deep neural faces show that synthetic faces offer a promising tool for both research and practical applications to help patients with deficits in speech perception.

## Acknowledgments

This research was supported by NIH R01NS065395 and U01NS113339.

